# Knowledge Mapping of Alternative Splicing of Cancer From 2012 to 2021: A Bibliometric Analysis

**DOI:** 10.1101/2022.10.12.511996

**Authors:** Bo Tian, Yan Bian, De-Jian Bian, Ye Gao, Xun Zhang, Si-Wei Zhou, Yan-Hui Zhang, Ya-Nan Pang, Zhao-Shen Li, Luo-Wei Wang

## Abstract

**Background:** As a processing method of RNA precursors, alternative splicing plays a critical role in normal cell physiological activities. Aberrant alternative splicing events are associated with cancer development and are promising targets for cancer treatment. However, no detailed and unbiased study describes the current state of alternative splicing of cancer research. This study aims to conduct a bibliometric analysis of alternative splicing of cancer research in the last decade to measure and recognize the current state and trends.

**Methods:** Web of Science Core Collection was used to acquire the articles. Utilizing three bibliometric tools (CiteSpace, VOSviewer, R-bibliometrix), we were able to measure and recognize the influence and collaboration data of individual articles, journals, and co-citations. Analysis of co-occurrence and bursts information helped us identify the trending research areas related to alternative splicing of cancer.

**Results:** From 2012 to 2021, the total number of papers on alternative splicing of cancer published in 766 academic journals was 3,507, authored by 20,406 researchers in 405 institutions from 80 countries/regions. Research involving alternative splicing of cancer genes was primarily conducted in the United States and China, with the United States retaining the prime status; simultaneously, the Chinese Academy of Sciences, Fudan University and National Cancer Institute were the institutions with strong research capabilities. Scorilas Andreas is the scholar with the most publications, while the most co-citations were generated by Wang, Eric T. Plos One published the most papers on alternative splicing of cancer, while J biol chem was the most co-cited academic journal in this field. Furthermore, these publications covered a wide range of areas, but were still primarily focused on the fields of molecular, biology, and immunology. The results of keyword co-occurrence analysis can be divided into three types: molecular (P53, CD44, Androgen receptor, SRSF3, ESRP1), pathological process (apoptosis, EMT, metastasis, angiogenesis, proliferation), and disease (breast cancer, colorectal cancer, prostate cancer, hepatocellular carcinoma, gastric cancer).

**Conclusion:** Research on alternative splicing of cancer has been a hot topic over the past decade, increasing in intensity each year. Current alternative splicing of cancer studies focused on hallmarks of alternative splicing in cancer, and alternative splicing signatures including diagnostic and therapeutic targets. Among them, the current trends are splicing factors regulating epithelial-mesenchymal transition and other hallmarks, aberrant splicing events in tumors, and further mechanisms. These might give researchers interested in this field a forward-looking perspective and inform further research.

## 1 INTRODUCTION

The progress of normal cell physiological activities depends on the spatiotemporal specific expression of genes, and abnormal expression often leads to pathological states including carcinogenesis(1). In the central dogma, mRNA is the pivotal molecule for the transmission of genetic information, and the stability of its structure and function in the process of transcription and translation is the basis for the smooth expression of genes and the production of functional proteins(2). Mature mRNA production needs to go through stages of RNA precursor processing and maturation, including splicing, adding cap structure, polyadenylation, and nucleobase modification… (3). These processing steps are regulated by their respective regulatory systems, further increasing transcriptome and proteome diversity.

Alternative splicing (AS) is one of the processing methods of RNA precursors, including the critical process of removing introns and connecting exons(4). Multiple transcripts can be transcribed from a gene to further enrich protein structure and protein domain activity(5). AS is completed by a synthetic spliceosome protein complex and is regulated by the interaction of cis-elements and trans-factors(6,7). This precise regulation maintain the balance of expression of different gene transcripts and ensure the accurate process of physiological activities and cell homeostasis(8). Due to abnormal regulation of cis-sequences and abnormal expression of splicing factors, abnormal splicing leads to the overproduction of tumor-promoting transcripts and the reduction of tumor-suppressing transcripts, which play an important role in the occurrence and development of tumor malignant phenotypes(9–11).

In recent years, research on the panoramic delineation and regulatory mechanisms of AS events in tumorigenesis has developed rapidly(4,7,12). Abnormal tumor-promoting transcripts, abnormally expressed splicing factors and mutated cis-element sequences can establish diagnostic models, and design small-molecule compound drugs or oligonucleotide therapeutic drugs in a targeted manner(13–16). These studies have expanded the research and development direction of new targets and specific drugs for early diagnosis and treatment of tumors.

Numerous reviews have summarized research on AS in cancer diagnosis and treatment, but to our knowledge, there is no comprehensive picture of AS splicing in cancer. Bibliometrics enables qualitative and quantitative analysis of data analysis, including contributions and collaborations of authors, institutions and countries, and assessment of research trends and themes(17–19). Thence, this report aimed to use the bibliometric method to assess the overall research trends of AS splicing in tumors and the hot issues over the past decade.

## 2 MATERIALS AND METHODS

### 2.1 Data Collection

Web of Science, which is broadly used in bibliometrics, can provide extensive and authoritative global academic data bibliometric software needs (20,21). We mainly use Web of Science Core Collection (WoSCC). The data were obtained on July 21, 2022, from the WoSCC database. The search formula was (TS= (alternative splicing)) AND TS= (cancer* OR anticancer* OR tumor* OR oncology OR neoplasm* OR carcinoma* OR lymphoma* OR sarcoma* OR leukemia*). The wildcard character (*) was used to allow variable endings of keywords so that to capture as much data as possible. There was a limitation on the publication year (2012-2021). The inclusion of the English-language literature was limited to original articles and reviews. We downloaded the search results as “Full Record and Cited References” and “Plain Text”. Following this, the files were renamed as “download_*.txt” in order to be able to be analyzed by CiteSpace software.

### 2.2 Data Analysis

We used three bibliometric tools, CiteSpace 6.1.R2 Basic (Chaomei Chen, 2006), R-bibliometrix 4.0.0 (Aria Massimo and Cuccurullo Corrado, 2017), VOSviewer 1.6.18 (Nees Jan van Eck and Ludo Waltman, 2010) to conduct bibliometric evaluation and visualization, and Microsoft Excel 2021 for statistics and plotting part of the figure. The first step was to clean our data. For example, “splicing factors” and “splicing factor” were merged as “splicing factor”, “rna splicing”, “pre-mrna splicing”, and “alternative splicing” were unified as “alternative splicing”(22).

CiteSpace is capable of discovering collaboration, keywords, domain research structure, future direction, and evolution as a bibliometric and visual analysis tool in a scientific field(23). Using CiteSpace, we utilize its co-occurrence, timeline, bursts, and dual-map functions to draw a series of figures about countries/regions and institutions, journals, references, citations, and keywords. We use the analysis process and parameter settings recommended by the software developers, where the time span is 2012-2021, the time slice is one year, the selection criterion is the g index (k = 25), pruning is none, and the minimum duration of burstness is two years. The size of the node in the CiteSpace visualization portrays the number of co-occurrence. Furthermore, linkages show the relationships between the co-occurrences. (24). As time passes from 2012 to 2021, the node and line’s colors change from purple to red to symbolize the different years. High betweenness centrality (≥0.10) nodes with purple circle serve as a hub between distinct networks (23,25,26).

Another bibliometric tool that excels in producing and visualizing knowledge maps is VOSviewer, which displays the kinds of clusters, overlays, or density colors (27). We mainly applied the co-occurrence analysis function of VOSviewer, including authors, journals, references, and their co-citations, as well as keywords. The specific parameters applied are mentioned in the corresponding chapters. In addition, we use the full counting method as counting method. The meaning of the size of the node is the same as that of citespace. Nodes with the same color belong to the same cluster. Additionally, links show the relationship between co-occurrences, and their degree of thickness is determined by the estimated strength value. The value is related to the number of papers published by the two authors or the frequency of the co-occurrence of the two keywords(27). The co-cited frequency is positively correlated with the word and round sizes as well as the yellow opacity in density maps. The color on the overlay map represents the typical publishing year.

R-bibliometrix is an R package for executing a comprehensive science mapping analysis of scientific literature(28). To be adaptable and make integration with other statistical and graphical R packages easier, R-bibliometrix was programmed in R. Maps of the geographical distribution of the countries/regions were produced using it. The network map displays the current state of research and communication between various countries/regions(28).

Excel 2021 software was used to anatomize the yearly publications and cited frequency. Additionally, we obtained impact factor (IF), journal citation report (JCR), average per item (ACI), journal classification, and author H-index from Web of Science on August 1, 2022.

## 3 RESULTS

### 3.1 Annual Growth Trend

From the WoSCC database, we acquired 3,544 papers, and we ultimately included 3,507 publications that qualified **(Figure 1**; **Supplementary Table 1**). The quantity of publications related to AS of cancer and the frequency of citations have both steadily increased over the past ten years, as illustrated in **Figure 2**.

**Figure 1.**
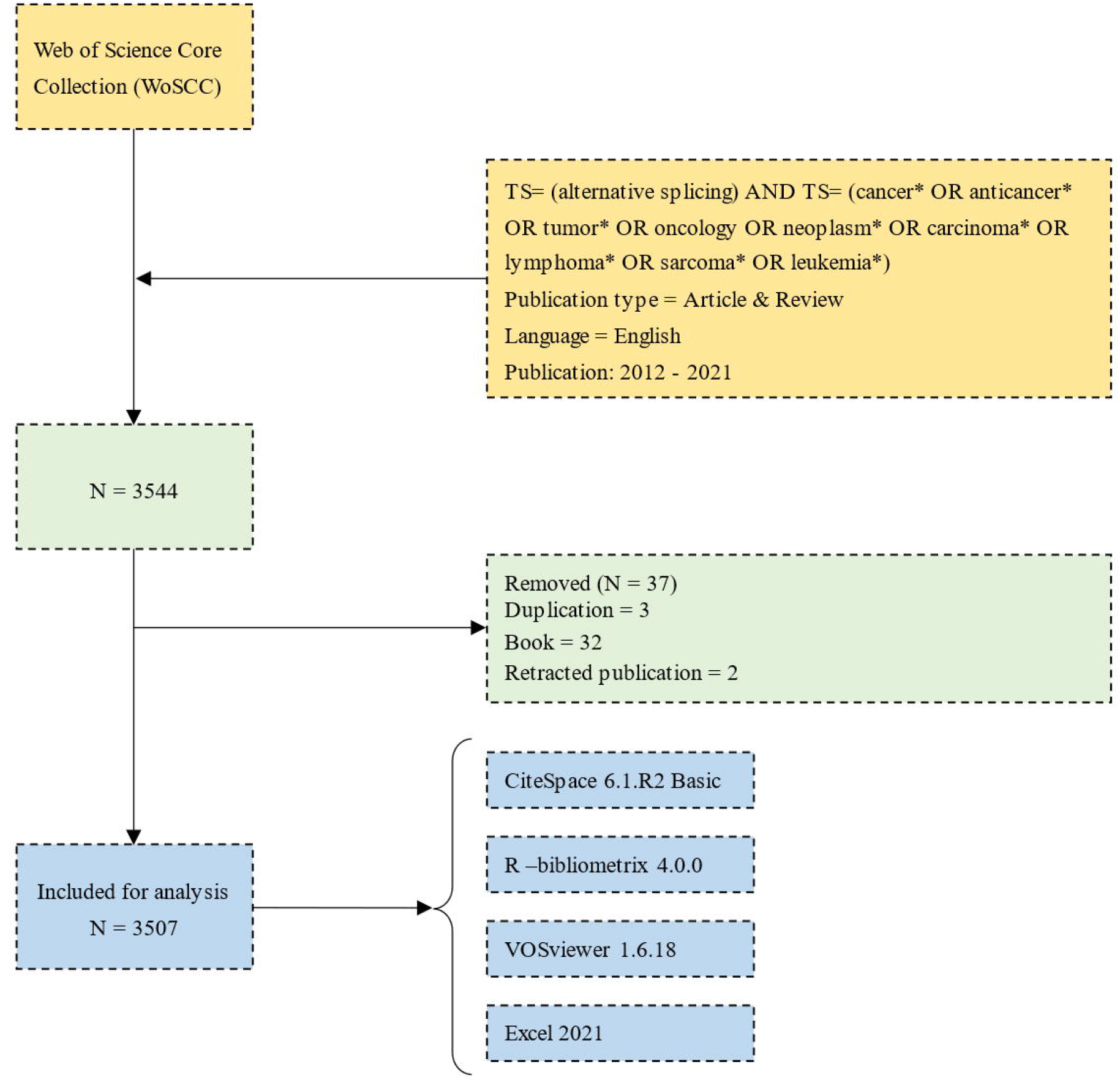
Flowchart of data collection, cleaning and analyzing.

**Figure 2.**
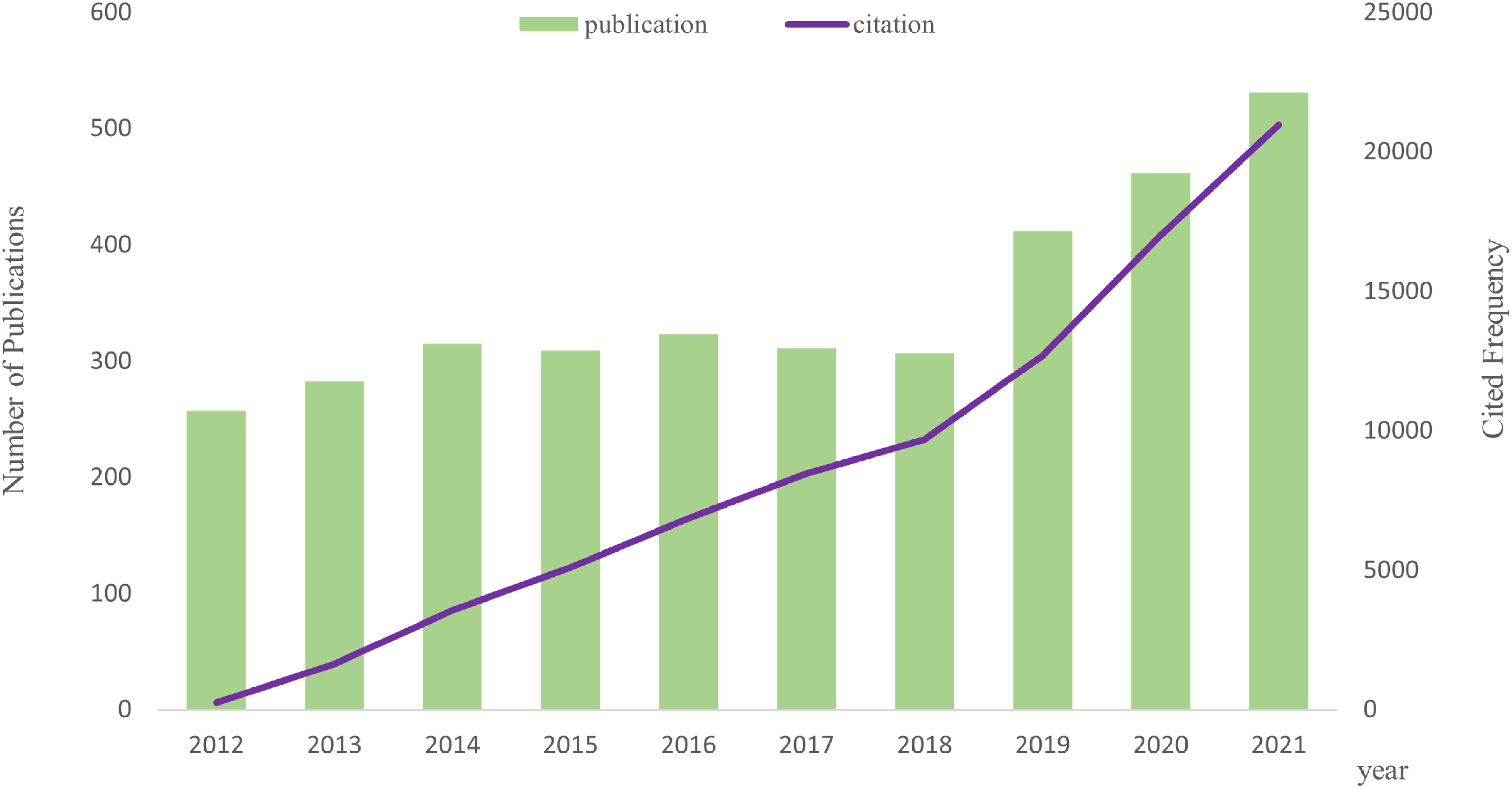
Tendency via bar and line graphs of alternative splicing of cancer publications and cited frequency nearly ten years.

### 3.2 Distribution of Countries/Regions and Institutions

3,507 papers total, representing 80 different countries/regions and 405 institutions. The United States, China, and Germany are the top 3 countries in terms of the number of published articles, with 1,165, 1,013, and 249, respectively (**Table 1**). China’s centrality, however, was less than 0.10, suggesting that it may not be a “hub” node in the AS of cancer studies(20). In contrast, the United States (centrality=0.41), England (centrality=0.25), and Germany (centrality=0.20) had high centrality, which is depicted in **Figure 3A** by a purple circle. As shown in **Figure 3A**, the closeness of countries/regions co-occurrence atlas was 0.1864, denoting lively collaboration between them(25). **Figure 3C** showed the relative proportion of annual publications for the top 10 countries from 2012 to 2021. It was evident that China’s share had gradually increased. A country/region co-authorship network was created by R-bibliometrix (**Figure 3D**). The network map showed the current state of research and communication activities between these countries/regions. The Chinese Academy of Science is the scientific research institution with the largest number of published papers, as shown in **Figure 3B**, yet its centrality is just 0.07 (n=76). By contrast, National Cancer Institute (n=65, centrality=0.14), University of California San Diego (n=48, centrality=0.2), and Karolinska Institutet (n=25, centrality=0.14) had a high centrality. The publication counts, h-index, and ACI of the top 10 most productive institutions were clearly displayed in the bar graph of **Figure 3E**.

**Table 1.**
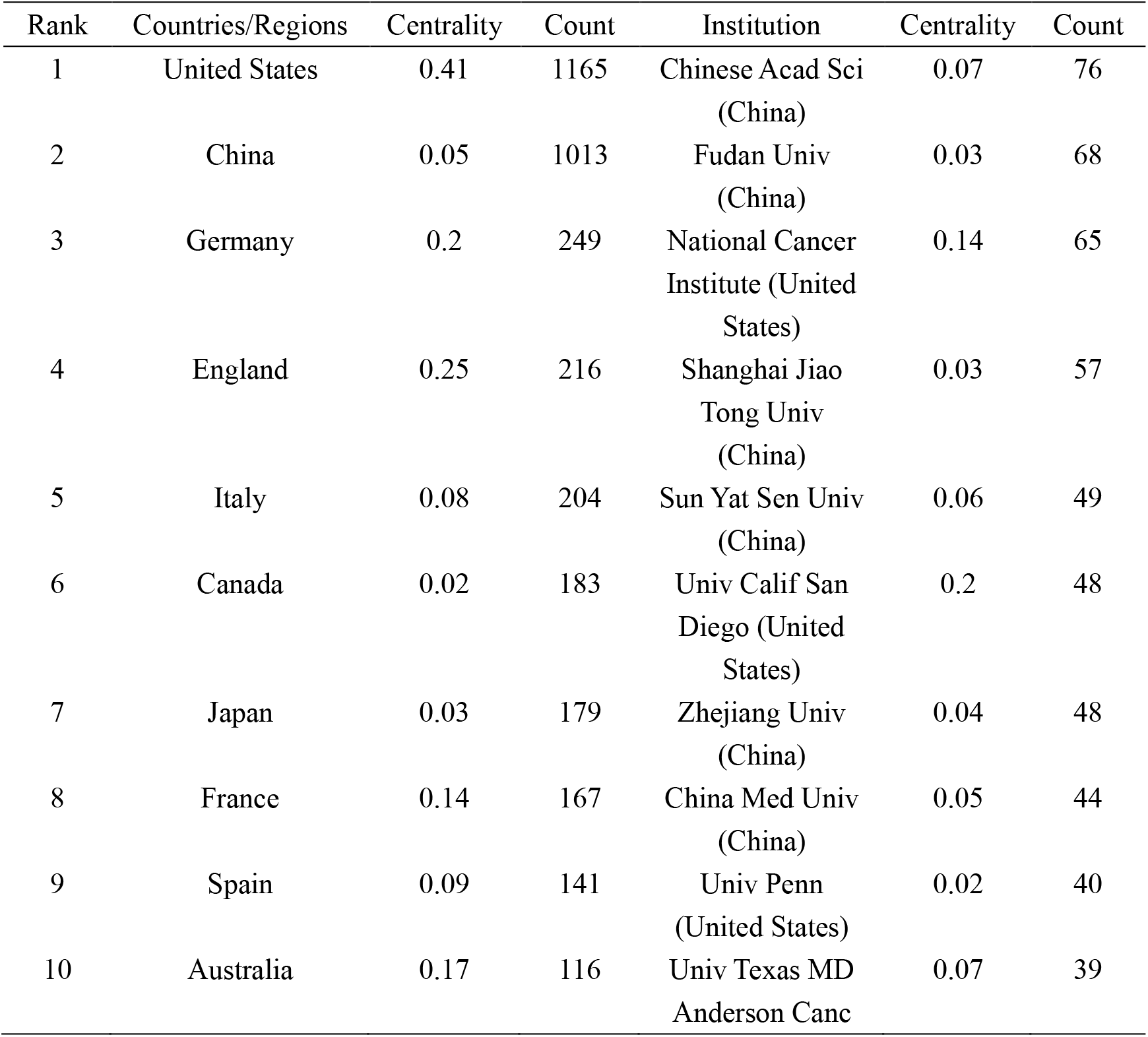
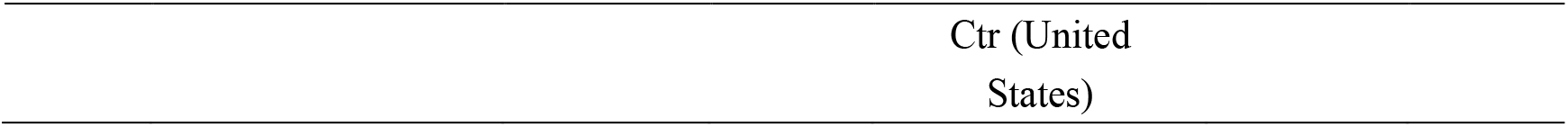
Top 10 countries/regions and academic institutions involved in alternative splicing of cancer research.

**Figure 3.**
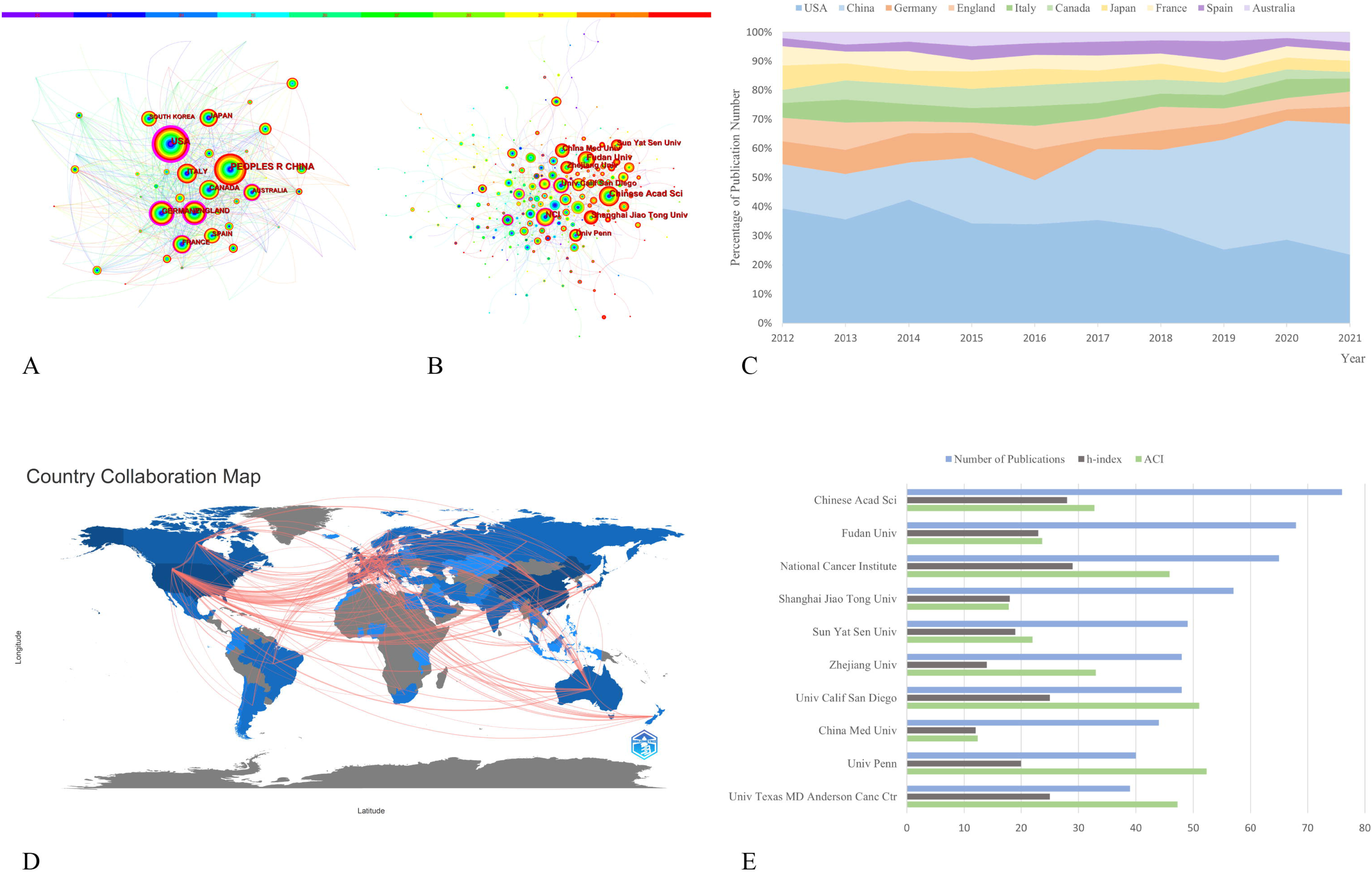
The co-occurrence atlas of (A) countries/regions (n≥100) and (B) academic institutions (n≥40) in alternative splicing of cancer research. As time passes from 2012 to 2021, the color of the node and line changes from purple to red. Purple-round nodes indicate strong betweenness centrality (≥0.1). (C) The relative fraction of annual publications in the top 10 countries from 2012 to 2021. (D) Network diagram of countries/regions (Min edges = 2) involved in alternative splicing of cancer research. (E) The top 10 most productive institutions’ publication counts, h-index, and ACI.

### 3.3 Authors and Co-Cited Authors

There were 20,406 authors active in AS of cancer research, and 25 of them published ten or more articles, as shown in **Figure 4A** and **Supplementary Table 2**. Scorilas Andreas, a National & Kapodistrian University of Athens scholar, was the most prolific author(n=25), followed by Adamopoulos Panagiotis G. and Karni Rotem (**Table 2**). Different colors represent different clusters in **Figure 4**, a total of 15 (29). Active partnerships, such as Ladomery Michael and Oltean Sebastian, are frequently found in the same cluster. Additionally, there were alliances between linked two authors of different colors, that seems to be Ghigna Claudia and Fu Xiangdong.

**Figure 4.**
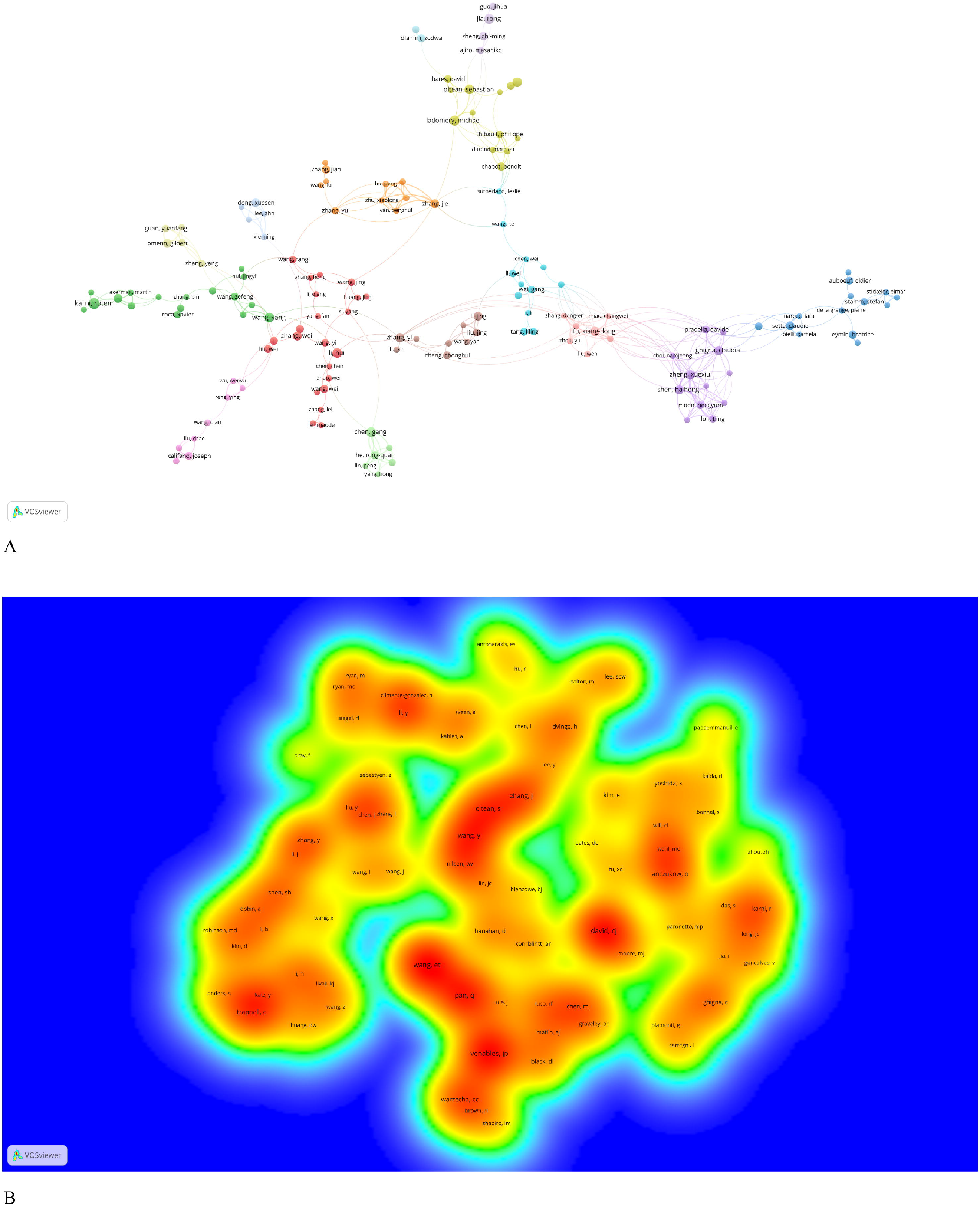
The co-occurrence (A) authors’ (documents ≥5) cluster map and (B) co-cited authors’ (citations ≥100) density map of alternative splicing of cancer research. (A) Nodes with the same color represent that they belong to the same cluster, the size of the node is proportional to the number of articles published by the author, and the thickness of the connection is proportional to the number of articles co-published by the two authors. (B) The co-cited frequency is positively correlated with the size and depth of the word and yellow, respectively.

**Table 2.**
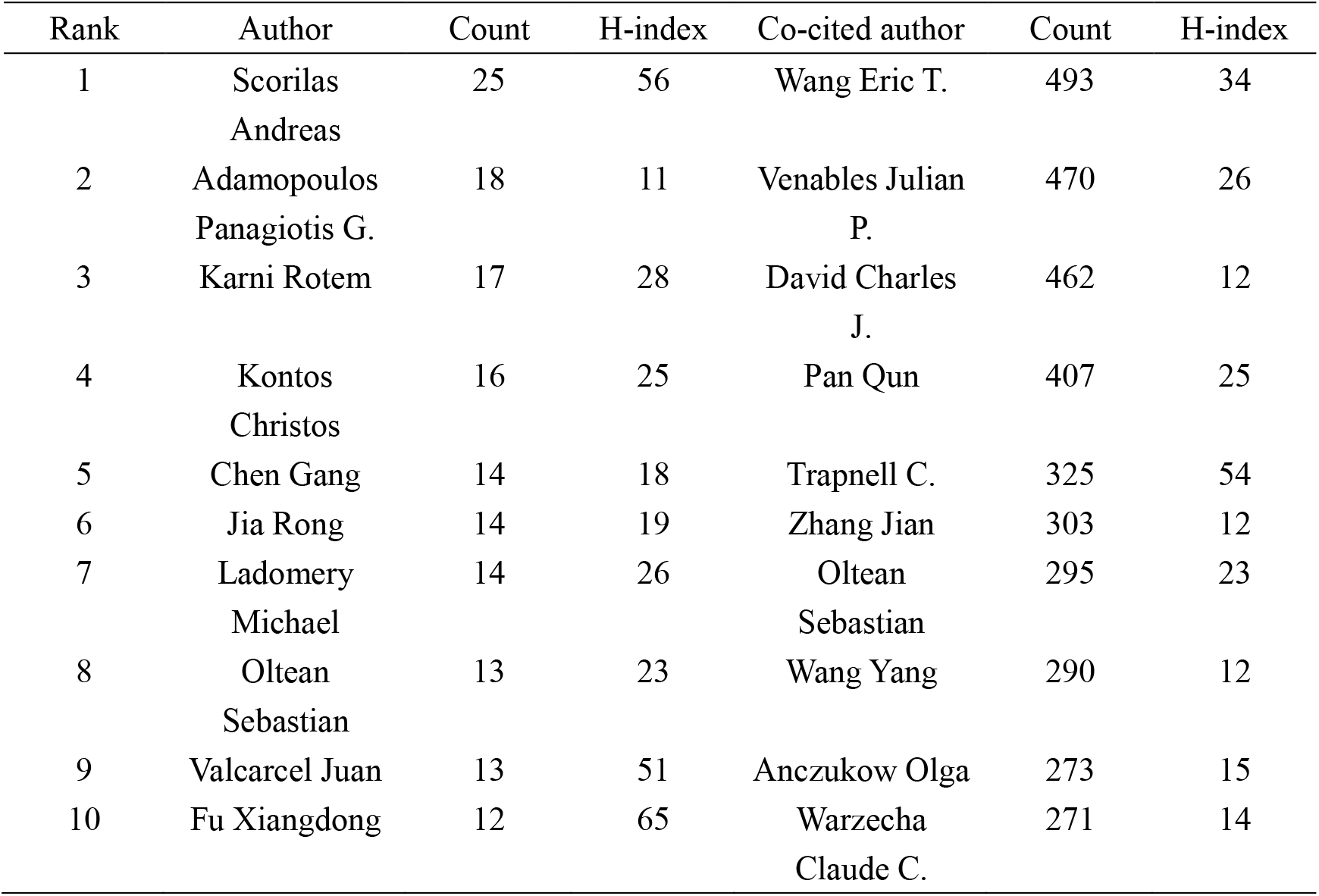
Top 10 authors and co-cited authors related to alternative splicing of cancer.

Authors who have been cited in one article are known as co-cited authors(30). We can see from **Figure 4B** and **Supplementary Table 2**, 83 authors out of the 85,667 co-cited authors had more than 100 co-citations. They were displayed as a density map in **Figure 4B**, which made it easy to identify the writers who were co-cited frequently. The hue gets warmer with more citations(31). Wang Eric T, Venables Julian P., and David Charles J. had the most co-citations, as indicated in **Table 2** and **Figure 4B**. The images cannot display all the information due to CiteSpace and VOSviewer visualization’s intrinsic restrictions. As a result, we have supplemented this with detailed information on the graphs in supplementary material.

### 3.4 Journals and Co-Cited Academic Journals

Articles about AS of cancer research have been published in 766 scholarly journals overall. Nine hundred five papers, or 25.81% of total publications, were published in the top 15 journals (**Table 3**). The number of papers published in Plos one is the largest (n=145), followed by the International Journal of Molecular Sciences (n=84) and Oncotarget (n=77). A co-citation map of journals was produced by VOSviewer, as shown in **Figure 5A**. The required minimum was established at 100 citations, and 292 journals satisfied the requirement.

**Table 3.**
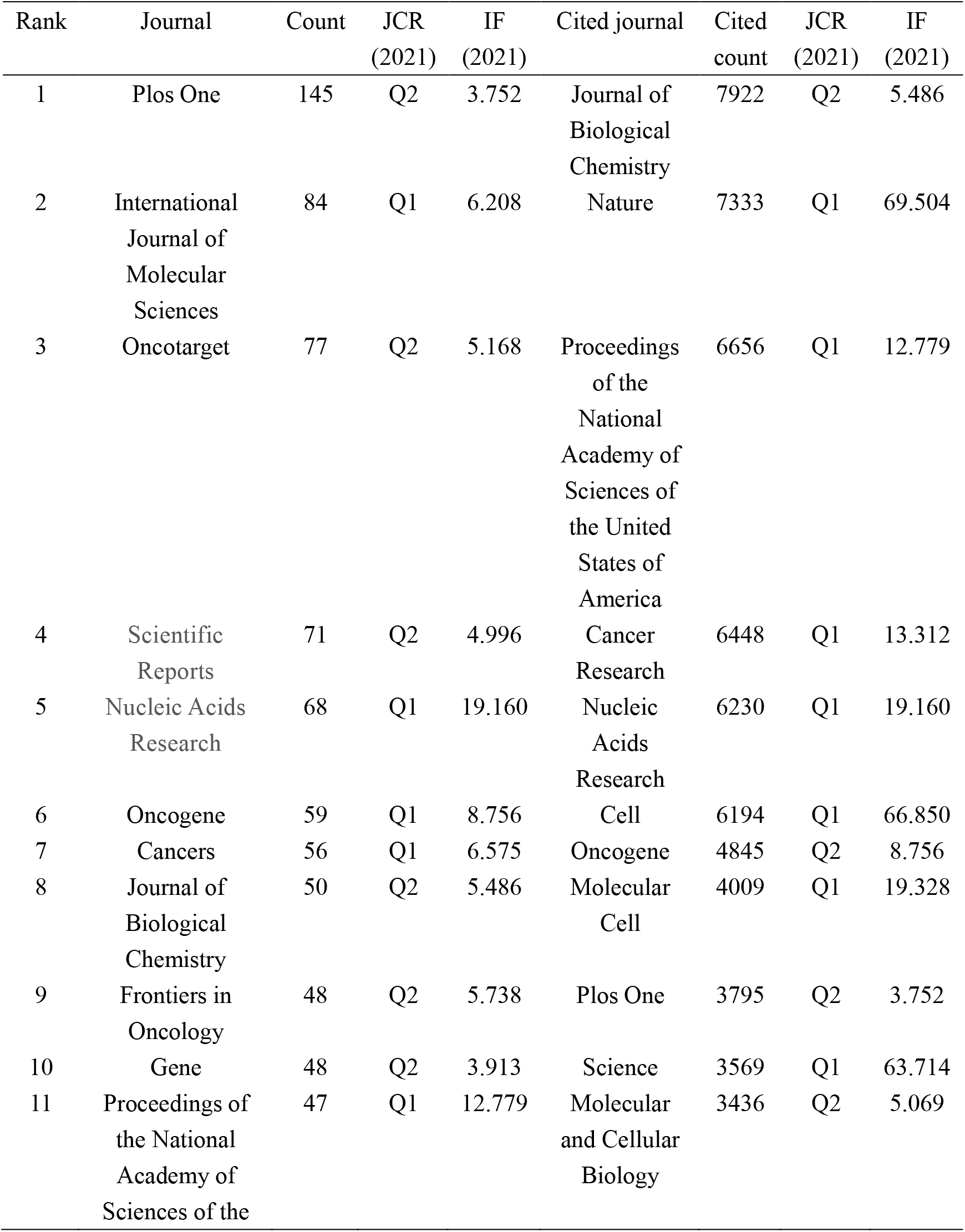
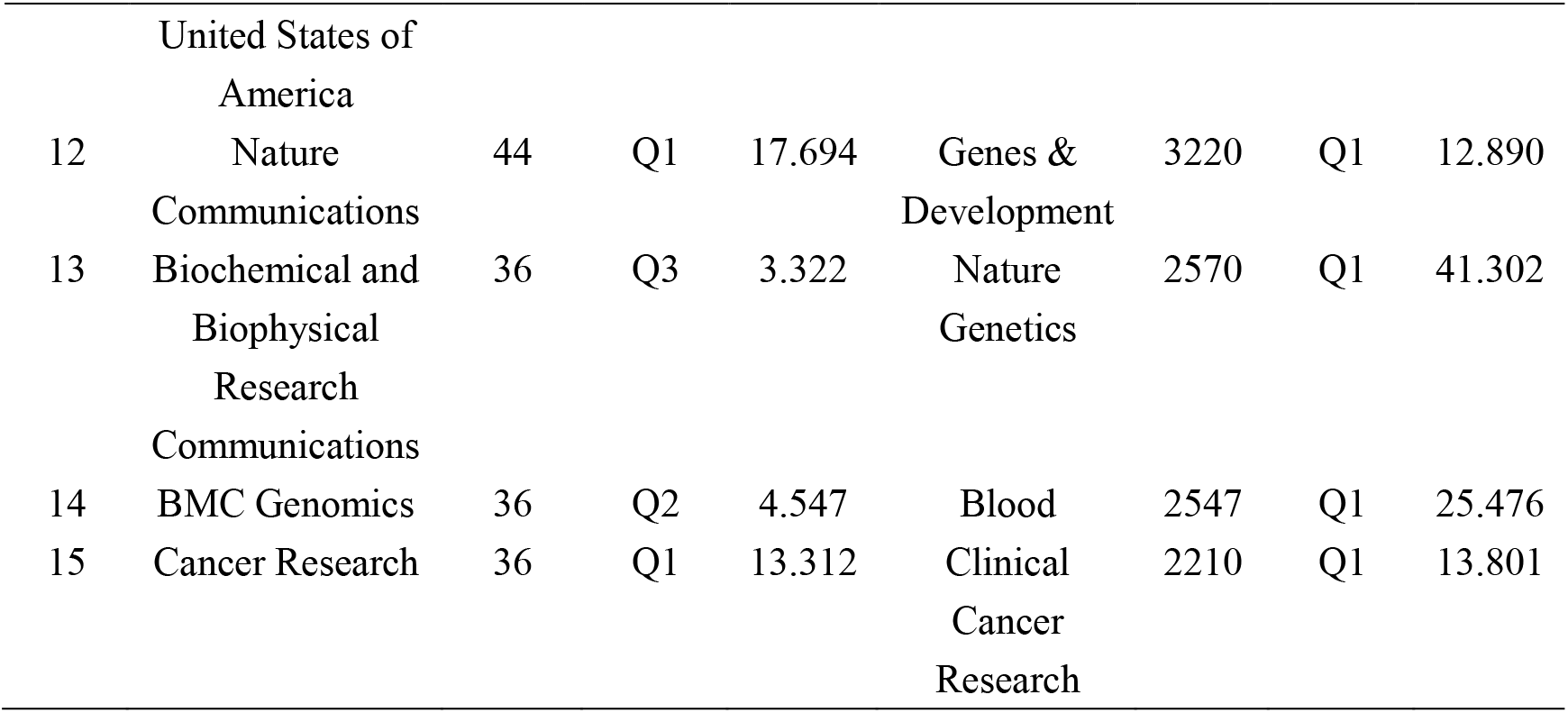
Top 15 journals and co-cited journals related to alternative splicing of cancer.

**Figure 5.**
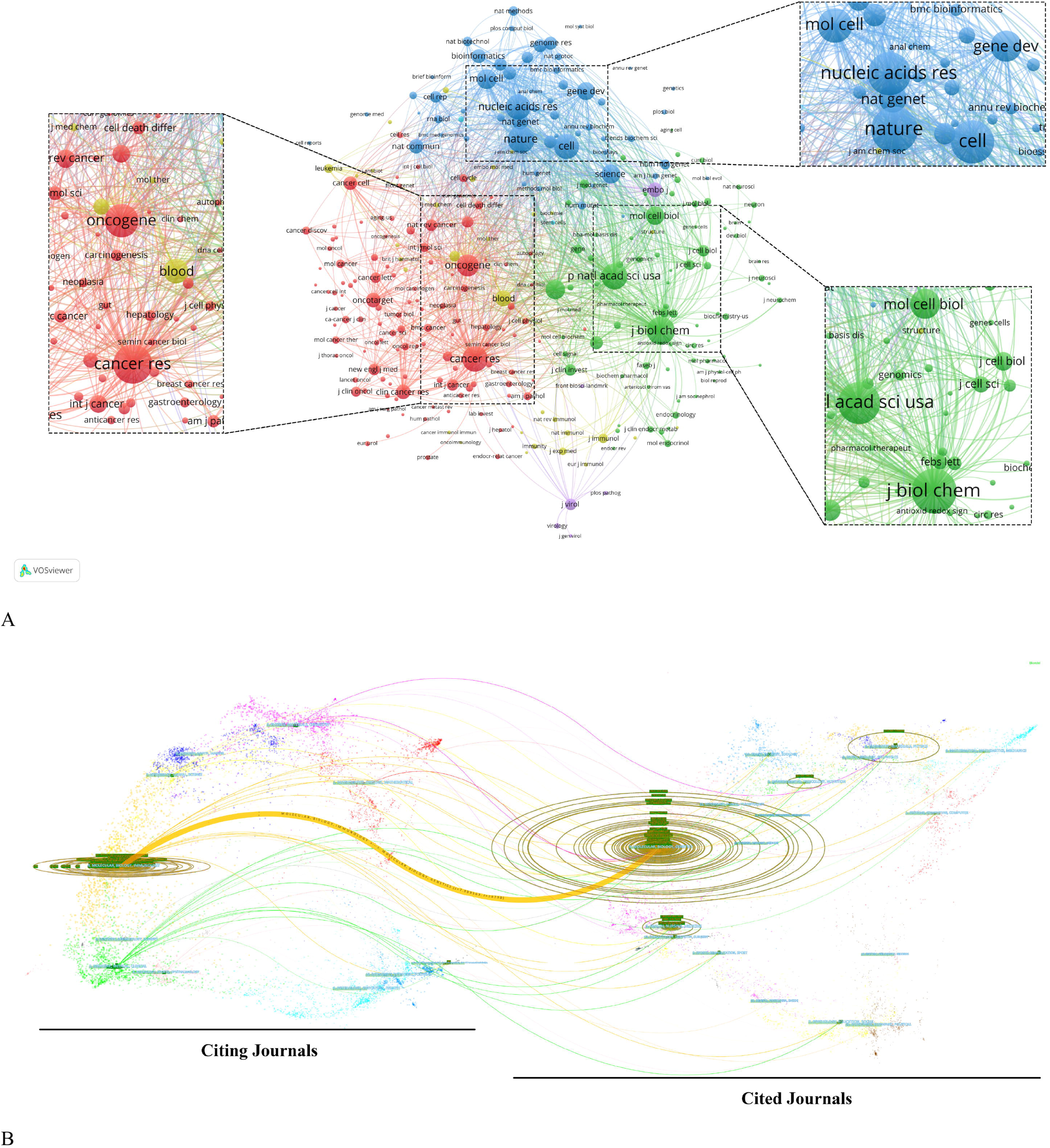
(A) Analysis of journals on alternative splicing of cancer co-citations via VOSviewer. (B) The dual-map overlay of journals on alternative splicing of cancer. The colored channel denotes the citation relationship, with the citing journals on the left and the cited journals on the right.

Of the 6,707 cited journals, 84 have more than 500 citations. Among them, Journal of Biological Chemistry (n=7,922), Nature (n=7,333), and Proceedings of the National Academy of Sciences of the United States of America (PNAS) (n=6,656) ranked in the top 3 in the number of citations (**Table 3**). Additionally, 31.16% of all cited sources came from the top 15 co-cited journals.

The dual-map overlay of journals represents the field distribution of citing and cited journals, as seen in **Figure 5B**(32). Citation relationships are indicated by colored paths, with citing journals on the left and cited journals on the right(24). The principal citation channel indicates that papers published in Molecular/Biological/Immunology journals mainly cite papers published in Molecular/Biological/Genetics journals (**Figure 5B**).

### 3.5 Co-Cited References and Reference Burst

37 references out of the 132,804 cited ones were quoted at least 100 times (**Supplementary Table 3**). The top 10 co-cited references were included in **Table 4**, with a minimum of 157 co-citations. The article by Wang Eric T. et al. from Nature in 2008 (n=465) is the one that has received the most co-citations out of all of them. In addition, four of the top 10 were reviews, and six of the top 10 were research articles.

**Table 4.**
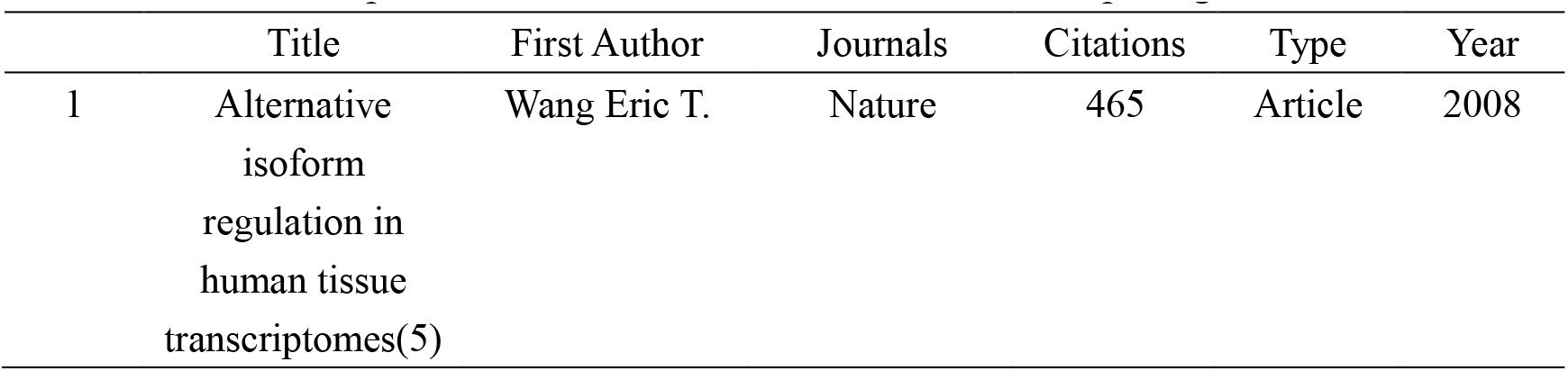
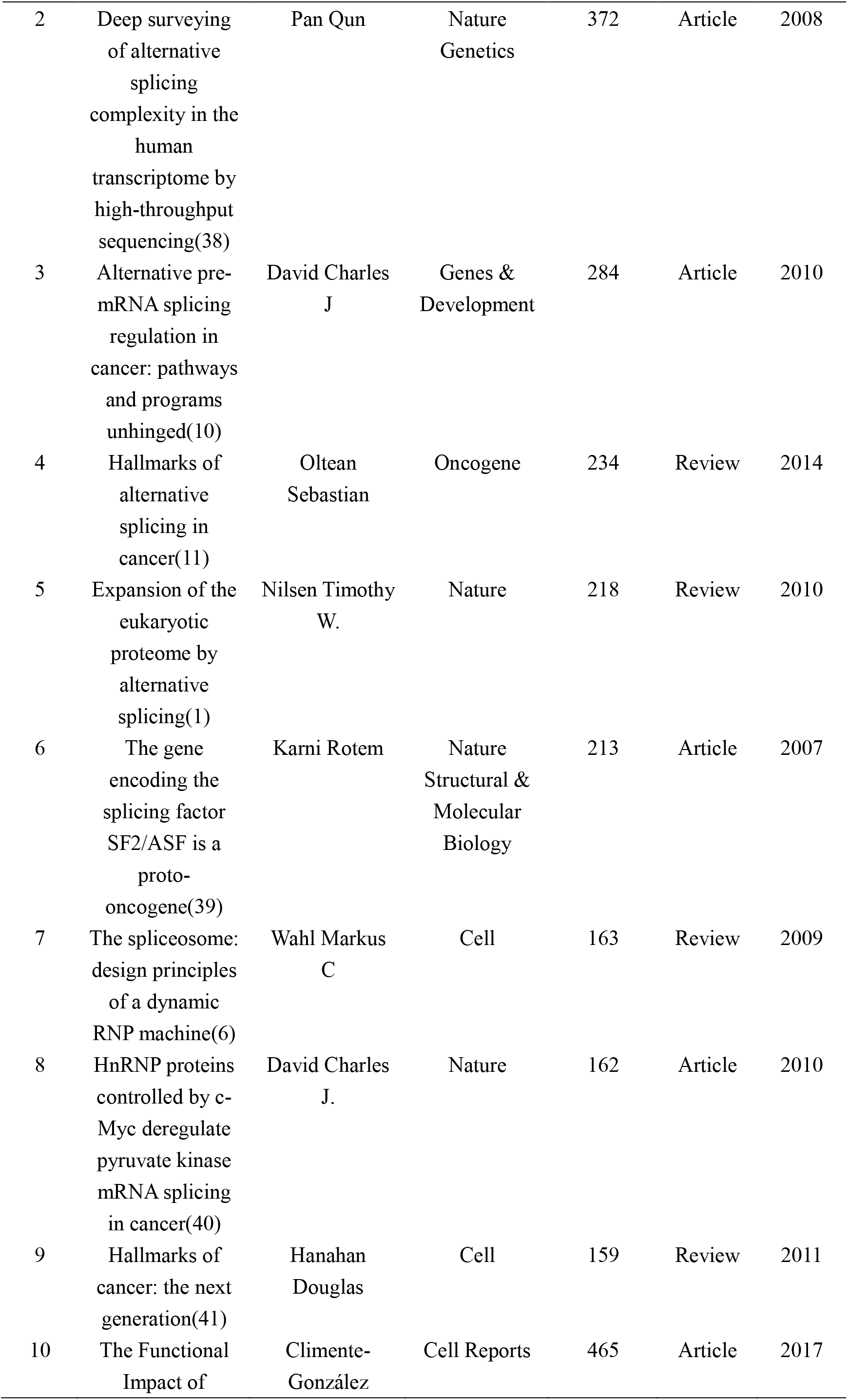
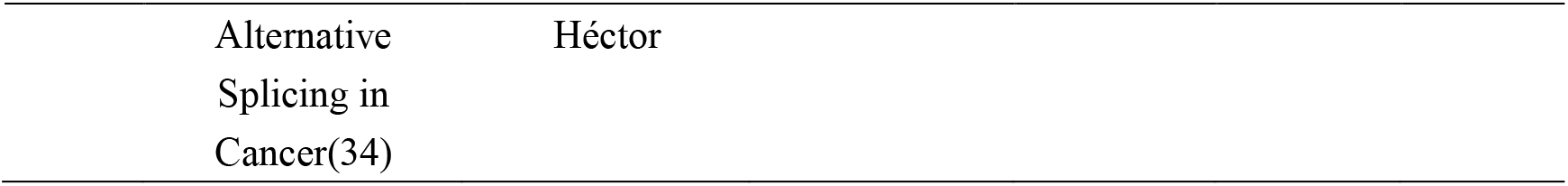
Top 10 co-cited references related to alternative splicing of cancer.

The entire network map might be partitioned into several clusters using the clustering function, and studies inside a cluster might have distinct study themes to studies from other clusters (**Figure 6A**). Each cluster’s most prevalent terms were designated as cluster labels(33). The references timeline view could allow users to see how various research hotspots have changed over time. As shown in **Figure 6B**, cluster #0 (post-transcriptional regulation), #3 (prognostic alternative), #4 (alternative splicing), #6 (circular rna), and #7 (androgen receptor), and #8 (noncoding rna malat1) started earlier; while cluster #1 (splicing signature), #2 (aberrant splicing) and #5 (oncogene srsf3) are still ongoing, which could be considered as the frontier.

**Figure 6.**
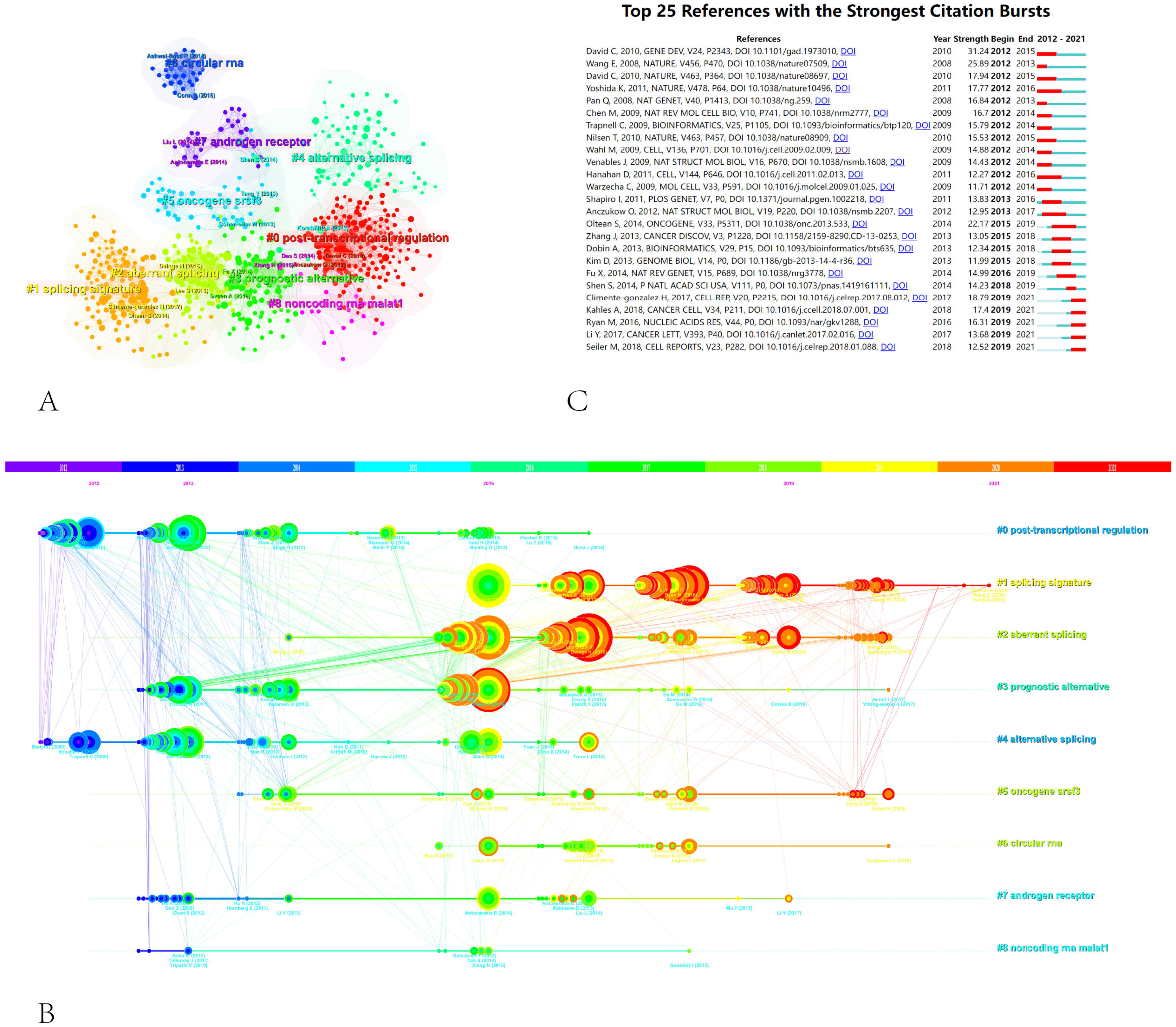
The reference co-citation analysis maps in (A) cluster view and (B) timeline view was produced by CiteSpace. (C) Visualization map of the top 25 references related to alternative splicing of cancer that has received the most citations. (B) Each cluster is represented as a horizontal axis; The larger the number of the cluster label, the smaller the cluster. The linkages show co-cited associations, and the node size represents co-citation frequencies. The node and line colors indicate distinct years. LLR used the title to extract cluster labels. (C) The red bars indicate citation burstness, whereas the blue bars indicate that the reference has been published.

References with citation bursts indicate that their citations have grown by leaps and bounds in a particular period(25). Two hundred fifty references were included in the citation bursts analysis, and top 25 were listed in **Figure 6C**. One paper, entitled “Alternative pre-mRNA splicing regulation in cancer: pathways and programs unhinged”, had the strongest burstness (strength=31.24) with citation burstness from 2012 to 2015(10), which published in Gene Dev by David Charles J. et al. in 2010. It is worth noting that five references (15,34–37) were still in burstness. Climente-González Héctor et al. (34)explored the functional influence of AS in cancer; Kahles André et al. (15)conducted a comprehensive analysis of AS across tumors; Ryan Michael et al. (35)provided a web-based resource for exploring the AS patterns of TCGA tumors; Li Yuan et al. (36)discovered a series of AS signatures in non-small cell lung cancer; Seiler Michael et al. (37)analyzed the functional consequences of somatic mutation of splicing factor genes, respectively.

### 3.6 Keyword Analysis of Trending Research Topic

6,116 keywords in total were recovered, 121 of which appeared at least ten times, and there were at least 30 times of 31 keywords. As represented in **Table 5**, alternative splicing (n=1,238), cancer (n=215) and prognosis (n=125) were the three most popular keywords. We divided the keywords into three categories: molecules, pathological processes, and diseases associated with AS of cancer, and listed the top 15 keywords in **Table 6**, respectively. Obviously, P53 (n=39), CD44 (n=35), androgen receptor (n=31), srsf3 (n=24), and esrp1 (n=16) were several molecular keywords with the highest frequency; apoptosis (n=94), EMT (n=61), metastasis (n=55), angiogenesis (n=40), proliferation (n=34), and epigenetics (n=23) were several pathological process keywords with the highest frequency; and breast cancer (n=115), colorectal cancer (n=82), prostate cancer (n=76), hepatocellular carcinoma (n=76), and gastric cancer (n=41) were several diseases keywords with the highest frequency in AS of cancer studies.

**Table 5.**
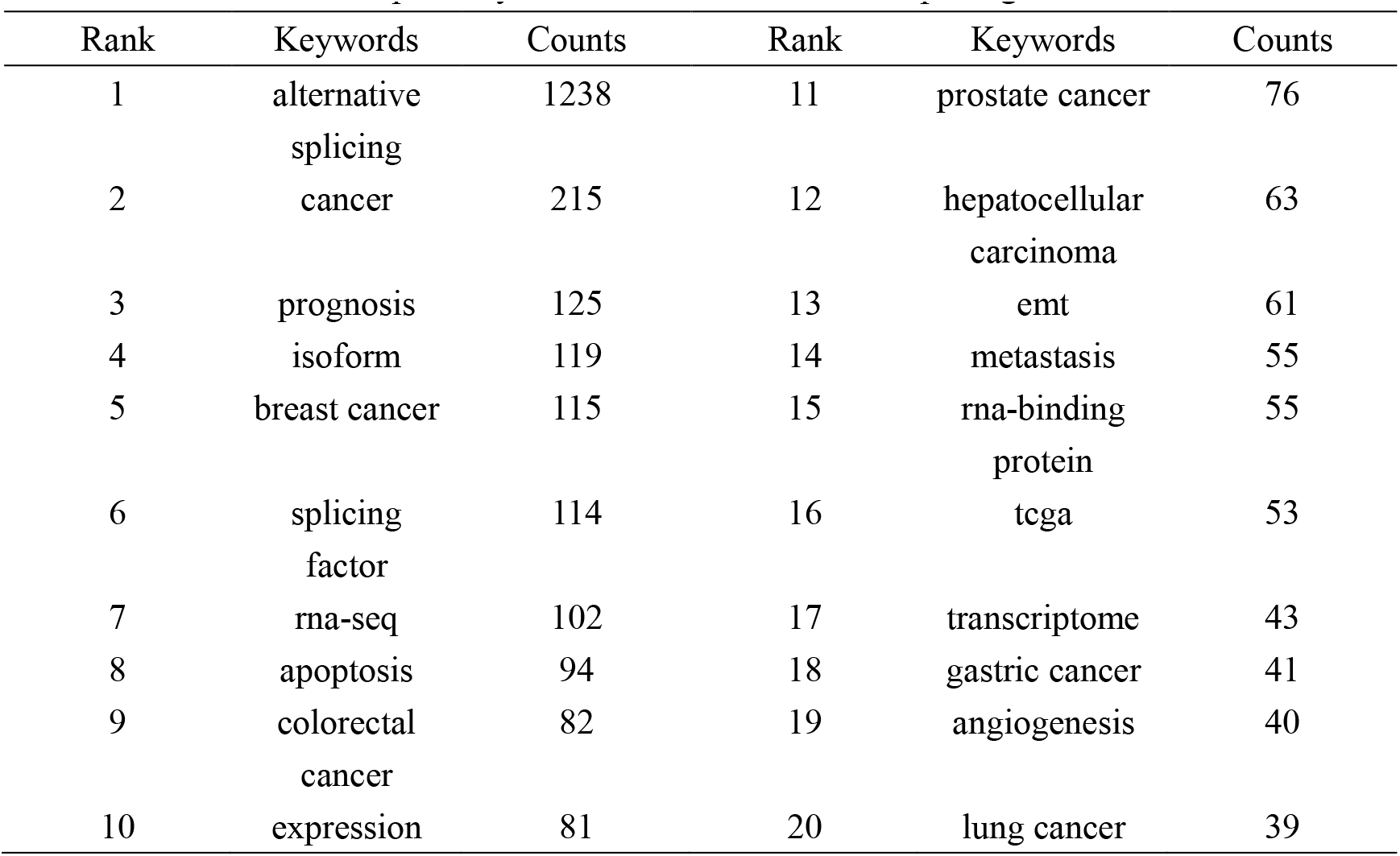
Top 20 keywords related to alternative splicing of cancer.

**Table 6.** Top 15 molecules, pathological process and disease related to alternative splicing of cancer.

| Rank | Molecules | Counts | Pathological process | Counts | Diseases | Counts |
| --- | --- | --- | --- | --- | --- | --- |
| 1 | P53 | 39 | apoptosis | 94 | breast cancer | 115 |
| 2 | cd44 | 35 | emt | 61 | colorectal cancer | 82 |
| 3 | androgen receptor | 31 | metastasis | 55 | prostate cancer | 76 |
| 4 | srsf3 | 24 | angiogenesis | 40 | hepatocellular carcinoma | 76 |
| 5 | esrp1 | 16 | proliferation | 34 | gastric cancer | 41 |
| 6 | srsf1 | 15 | epigenetics | 23 | lung cancer | 39 |
| 7 | brca1 | 14 | drug resistance | 22 | glioblastoma | 25 |
| 8 | rbm10 | 14 | invasion | 22 | non-small cell lung cancer | 25 |
| 9 | rbm5 | 14 | survival | 22 | pancreatic cancer | 25 |
| 10 | htert | 12 | tumor microenvironment | 20 | melanoma | 22 |
| 11 | mdm2 | 11 | autophagy | 19 | acute myeloid leukemia | 22 |
| 12 | ptbp1 | 11 | migration | 19 | glioma | 21 |
| 13 | srpk1 | 11 | metabolism | 18 | ovarian cancer | 21 |
| 14 | bcl-x | 10 | proliferation | 18 | cervical cancer | 18 |
| 15 | pkm2 | 10 | dna damage | 15 | lung adenocarcinoma | 16 |

Keywords with high co-occurrence counts (n≥10) were displayed as an overlay map in **Figure 7A**, with the hue denoting the typical year of publication. As we can see, the emerging fields that were given the color yellow include splicing factor, prognosis, immunotherapy, and TCGA (The Cancer Genome Alta). Each cluster displayed the top 3 keywords over time in the timeline view (**Figure 7B**). Six of the seven clusters were still active except for cluster #6. Among them, #0 (splicing event) is the biggest cluster, followed by #1 (therapeutic target), #2 (epithelial-mesenchymal transition), and #3 (splicing factor). **Supplementary Table 2** provided further details.

**Figure 7.**
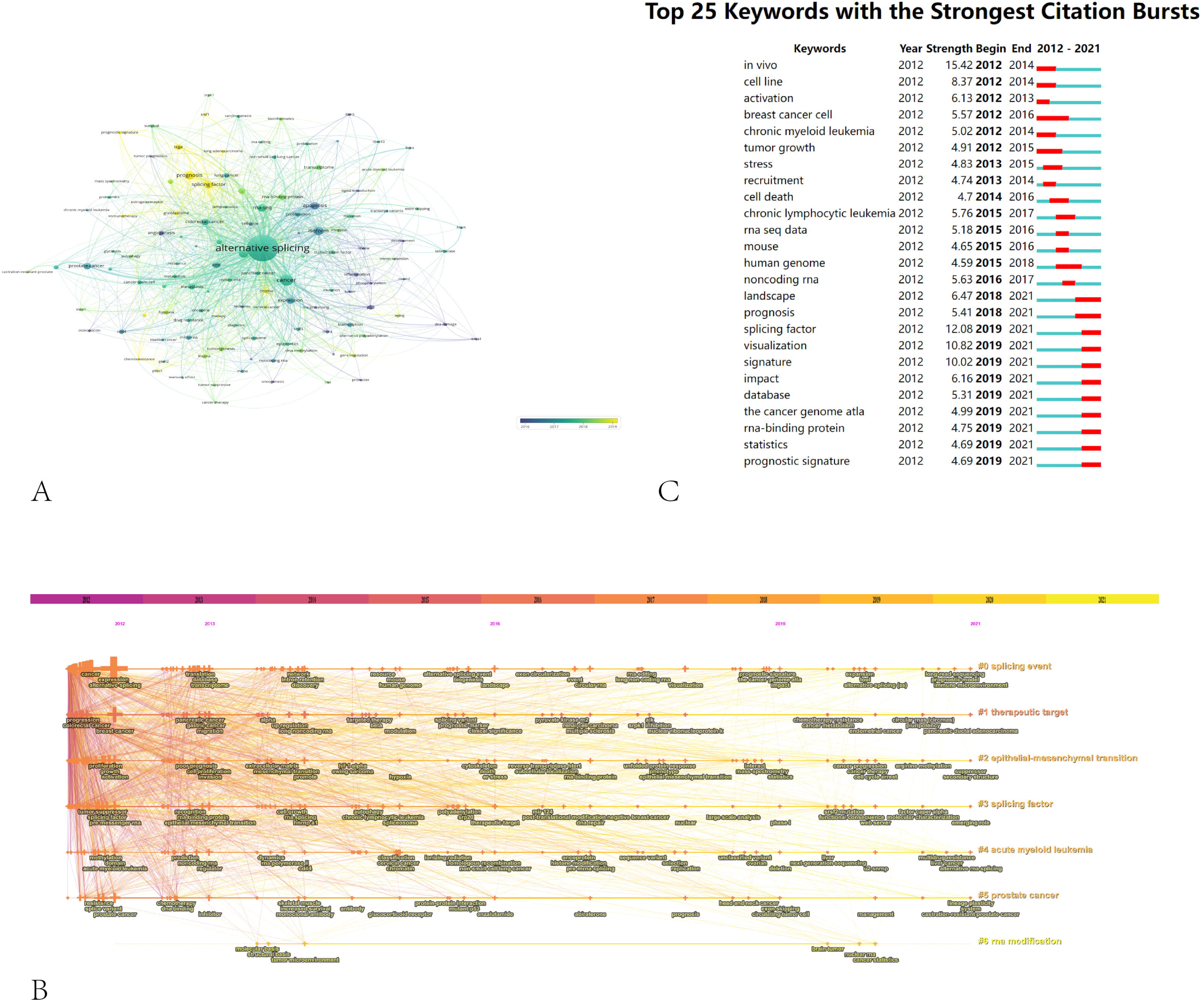
The (A) overlay map (n≥10, max lines = 1000) and (B) timeline view of keywords associated with alternative splicing of cancer. (C) Greatest citation bursts for the top 25 keywords (sorted by the starting year). (A) The color of the node represents the average publication year, the size of the node is proportional to the counts of co-occurrences of keyword, and the thickness of the link is positively correlated with the number of co-occurrences of the keywords represented by the two nodes (B) Each cluster is represented as a horizontal axis; The larger the number of the cluster label, the smaller the cluster. LLR used the title to extract cluster labels. (C) The red bars indicate citation burstness.

Bursts of keywords are those that were used a lot during a certain time frame(25). **Figure 7C** demonstrates that in vivo had the strongest bursts, with a strength of 15.42, followed by splicing factor at 12.08 and visualization at 10.82. Notably, until 2021, the landscape, prognosis, splicing factor, and other keywords were in burstness. The results of VOSviewer and CiteSpace had considerable some similarity regarding keywords, indicating the reliability of our analysis.

## 4 DISCUSSION

### 4.1 General Information

Based on information from the WoSCC database, a total of 3,507 AS of cancer publications by 20,406 writers in 405 institutions from 80 countries/regions were published between 2012 and 2021 in 766 academic journals. According to a rising trend in publications, curiosity and attention are growing to the AS of cancer. The AS research officially started in 1977, when Roberts R.J. and Sharp P.A. found the phenomenon of “alternative splicing”(42,43). Since that time, AS research has expanded quickly. Our focus is on the relationship between alternative splicing and cancer. AS research of cancer has steadily increased in steps during the last ten years, and the relevant published article and cited frequence in 2021 is almost twice and over eighty times that of 2012, respectively. The volume of published articles is a very important indicator, but the centrality is a crucial indicator for the quality of published articles in country/region analysis, where high centrality nodes (≥0.10) indicate the “hub” influence of particular countries/regions in the worldwide collaboration map(23,26,33,44). The United States and China made the largest contributions to papers on AS of cancer, as shown in **Table 1** and **Figure 3A**. The top 10 institutions by the number of publications were mainly from China and the United States, six were from China and four were from the United States, respectively. China and Chinese institutions, on the other hand, had a centrality of less than 0.1 while the United States had a centrality of 0.41, suggesting that the United States may continue to dominate AS of cancer field. Furthermore, the United States, England, Germany, Australia and France had high betweenness centrality, which indicates that they were important hubs in AS of cancer field’s international collaboration. Moreover, it could be seen that countries/regions and institutions had active collaboration in terms of network density, respectively.

Scorilas Andreas not only published the most articles related to AS of cancer but is also the top 1 H-index scholars **(Table 2)**, demonstrating his excellent contribution to AS of cancer research. Scorilas is a researcher at National & Kapodistrian University of Athens, committed to AS, gene transcription, next-generation sequencing. Most recent 10 years, his group published a series of articles(45–50) that described the process of identification of novel alternative-splicing variants using next-generation sequencing methodology and discussed their expression situation and pathophysiological implications. Besides, Oltean Sebastian is also a schalors working in University of Exeter; his team centered on regulation and form of AS in prostate cancer and therapeutic targets associated with AS in cancer. In 2014, Nature published an Oltean and Bates’s review entitled “Hallmarks of alternative splicing in cancer”(11). They summarized how the numerous phenotypic traits that tumors acquire are influenced by AS and identified a new class of anticancer treatments called alternative splicing inhibitors; This article co-cited up to 234 times with citation bursts from 2015 to 2019. Significantly, the top 1 co-cited author, Wang Eric T., a professor at University of Florida, focused on exploring the function of the short and long gamma subunit splice variants of human GABA(A) receptors. He published the top 1 co-cited reference that reviewed the regulation of AS(5), with the second strongest citation bursts strength.

According to a journal analysis **(Table 3)**, Plos One was the ninth most referenced journal and published the most AS of cancer studies. The fact that Nucleic Acids Research was among the top 5 academic journals as well as co-cited journals shows how important it is to the dissemination AS of cancer research. The top 10 most-cited papers were published in the journals, which are also generally the most-cited journals. For instance, Nature obtained the second-highest number of co-citations, in part due to three of the top ten extremely co-cited references(1,5,40)**(Table 4)**. Similar to the dual-map study **(Figure 5)**, we found that the majority of the journals were in the subjects of molecular, genetics, and comprehensive fields.

The knowledge network may be represented in part by the co-cited references cited by the papers in the corresponding field(24,25,51). The study’s top 10 co-cited articles found that, three articles primarily explored the relationship between AS and hallmarks of cancer(11,34,41), two were about the discovery of alternative-splicing isoforms using high-throughput sequencing(5,38), two were related to the regulation of AS(6,10), two papers focused on several specific molecules, splicing factor SF2/ASF and heterogeneous nuclear ribonucleoproteins (hnRNPs) controlled by c-Myc, respectively(39,40). And one review discussed the mechanisms of proteome expansion by AS(1). As shown in our citation bursts results (**Figure 6C**), five references in the AS of cancer research are still active: three are related to the function and mechanisms of AS(34), one is dedicated effort to mine prognostic AS signatures(36), and three present the landscape of AS in TCGA tumors(15,35,37).

### 4.2 The Hotspots and Trending

#### 4.2.1 Hallmarks of Alternative Splicing in Cancer

As carcinogenesis progresses, the transformation of the AS state occurs concurrently with the acquisition of cancer characteristics(11). Numerous tumor-specific isoforms are produced by erroneous splicing states, and the disordered expression of these isoforms propels the development of tumor malignant characteristics(52). Alterations in isoform-specific splicing patterns of many genes drive tumor cells to acquire sustained proliferative signals, escape growth inhibition, resist cell death, induce angiogenesis, obtain the ability to invade and metastasize, and achieve immune evasion mechanisms(4). As shown in **Figure 6C**, the results of the reference burst showed a review by Oltean Sebastian systematically expounded the hallmarks of AS in cancer. Among the key words of pathological process **(Table 6)**, it also covered many hot words such as apoptosis, migration, angiogenesis, and proliferation, which were closely related to cancer hallmarks.

In our study, the keywords co-occurrence analysis by VOSviewer (**Figure 7A**) and cluster label (#2) from CiteSpace (**Figure 7B**) both highlighted epithelial-mesenchymal transition (EMT), through which relatively quiescent, tightly connected epithelial cells acquire highly motile and invasive mesenchymal properties(53,54). Tumor cells develop an aggressive and migrating character throughout the malignant evolution of epithelial cancers, allowing them to invade nearby tissues and disseminate to distant organs(55). 90% of cancer deaths are caused by this metastasis development process(56). EMT is linked to the reprogramming of multiple genes’ expression. E-cadherin, claudins, and occludins are examples of epithelial-specific genes that are inhibited by the SNAIL proteins (SNAIL1 and SNAIL2)(57,58). N-cadherin, fibronectin, and matrix metalloproteases are examples of mesenchymal-specific genes that can be stimulated by the bHLH transcription factors (TWIST1 and TWIST2) and ZEB proteins (ZEB1 and ZEB2)(59–61). There is a ton of evidence to support the idea that AS events cause mesenchymal and epithelial cells to differ proteomically(62). According to reports, the modulation of a number of splicing factors is crucial to the EMT process(63). Numerous pre-mRNA targets can be regulated by a single AS factor. As a result, variations in their expression levels may have an impact on multiple aspects of the development of EMT(53).

The fifth keyword of Molecules in our study is ESRP1 (**Table 6**). Depending on where their binding sites (UGG-rich motifs) are located in their RNA targets, ESRP proteins have a positional effect and either promote or repress exon inclusion(64,65). ESRP1 and ESRP2 are two epithelial-restricted splicing regulators(66). During the activation of EMT programs, ESRPs regulate a network of epithelial regulators, and AS has a significant impact on the physical connections between isoforms(67). The isomers may have completely different effects. By boosting P120’s affinity for E-cadherin, P120 isoforms 3 and 4 can help epithelial cells adhere to one another(68). In contrast, p120 isoform 1 promotes RAC1 activity and stimulates cell migration and invasiveness by blocking the RHOA-ROCK signaling pathway(69).

Another molecule in **Table 6** closely related to EMT is CD44. Various extracellular matrix elements are bound by the cell surface glycoprotein that CD44 encodes for(70). Mesenchymal CD44 splicing isoforms can be produced more readily when ESRP1 is inhibited by ZEB1(71). Notably, the change from the epithelial isoform (CD44v) to CD44s reveals a crucial function in EMT(72).

The vast majority of genes are alternatively spliced, and numerous isoforms have been particularly linked to cancer hallmarks(11). Targeting specific phenotypes to identify different abnormal regulatory mechanisms in tumor development and metastasis is conducive to developing specific targeted drugs to inhibit cancer phenotypes development(73–75).

#### 4.2.2 Alternative Splicing Signature of Cancer

In our research, signature-related words appeared many times. For example, one of the cluster labels of the reference is splicing signature (#1, **Figure 6B**), therapeutic target (#1, **Figure 7B**) is in the keywords cluster labels, and signature and prognostic signature are in the keywords burst (**Figure 7C**). In the last ten years, transcriptome sequencing at the genome-wide level has demonstrated that AS is closely modulated in a tissue- and developmental stage-particular way, and it has also been shown that AS is frequently dysregulated in a variety of human cancer types(63,76–79). As a result, the study of AS events as tumor indicators and therapeutic targets has gained a lot of attention. Additionally, the range of tumor biomarkers and therapeutic targets has been considerably broadened by AS(80).

Currently, it is understood that the principal causes of tumorigenesis are splicing abnormalities, which include genetic changes in the spliced gene and changed expression of either or both of the key regulators or core components of the precursor messenger RNA (pre-mRNA) splicing machinery(81,82). This also provides a theoretical basis for AS to become a tumor marker and a therapeutic target.

The core issues in oncology continue to be the early detection and diagnosis of cancer as well as the selection of the best-customized treatment for each patient. It is now possible to identify genome-wide AS thanks to the advancement of high-throughput sequencing methods, particularly RNA sequencing(83–88). Numerous potential benefits of RNA-seq include its capacity to estimate the great amount both acknowledged and original alternative transcripts, as well as its ability to offer a finer resolution, deeper coverage, and greater accuracy(88). Due to advancements in sequencing and bioinformatics technology, a number of cancer-specific AS events with potential prognostic and predictive significance in clinical situations have been found thus far(36,89–93). For instance, hormone-directed therapy has been shown to be less successful in castration-resistant prostate cancer patients who carry the alternatively spliced androgen receptor variation 7 (94). It was proposed that the tumors with increased background expression of PKM2 show a more aggressive phenotype and poor response to chemotherapy in pancreatic ductal adenocarcinoma patients undergoing radical surgery and adjuvant chemotherapy(95). A prospective therapeutic target in colorectal cancer is CD44 variation 6, an independent negative prognostic factor(96–98). Similar to our research, these molecules appear in the molecular keywords (**Table 6**), indicating that they have potential clinical translation value and are research hotspots.

Numerous effective therapeutic approaches have been developed as a result of the discovery of cancer-specific AS mutations. First, reversing faulty RNA splicing has been made possible by inhibiting post-translational modifications of splicing factors or RNA binding proteins, particularly with the help of small-molecule inhibitors that target protein kinases(99,100). The dual-specificity Cdc2-like kinases and SR-rich protein-specific kinases are these agents’ two primary targets(101). Second, a critical treatment focus is the adjustment of signaling pathways that control AS events(102). For instance, the PI3K/AKT/mTOR pathway inhibitors MK2206 and BEZ235 can alter splicing results(103,104). Third, antisense oligonucleotides can hinder the splicing machinery’s ability to reach the regulatory regions in the pre-mRNA for therapeutic purposes and encourage the purge of the targeted mRNA by endogenous cellular nucleases(105,106). Bcl-x is a critical gene, ranked 14th in our molecular keywords (**Table 6**). Bcl-x antisense oligonucleotides were created to encourage a splicing transition that favors the generation of pro-apoptotic Bcl-xS rather than anti-apoptotic Bcl-xL(107). Fourth, AS isoform proteins unique to tumors have always been prospective therapeutic targets. Certain tactics have been devised to use immunotherapies to target cancer-specific isoforms(80). The EGFR isoforms de4 and vIII are among the most extensively researched therapeutic targets(108).

In the realm of tumor research, the identification and therapy of cancers are enduring focus topics. More significant biomarkers may be represented by AS signature spectra or characteristics. These AS signatures can be used as biomarkers for tumor diagnosis or to develop more effective drug candidates.

#### 4.2.3 Mechanism of Alternative Splicing

As shown in **Table 6**, it can be seen that splicing-related proteins such as srsf3, esrp1, srsf1, ptbp1, rbm10, and rbm5 have constantly been research hotspots. Among them, srsf3(#5, **Figure 6B**) ranks sixth in the reference cluster label, and the splicing factor (#3, **Figure 7B**) is also one of the keyword cluster labels. The keyword splicing factor is also reflected in the keywords burst (strength=12.08, **Figure 7C**), with burstness until 2021. This part mainly discusses the important role of splicing factors in AS. Splicing factors are auxiliary proteins that take part in the splicing of pre-mRNA. Trans-acting splicing factors, which bind to sequence motifs connected to the stimulation (enhancers) or inhibition (silencers) of splicing, usually control AS. These motifs can be found in exons and introns, and they frequently have the greatest impact near splice sites(109,110).

Our research indicates that splicing factors from the families of hnRNPs and serine/arginine-rich proteins (SR proteins) have been a focus of this field’s research. Many splicing factors ranked high in the molecular keywords (**Table 6**) belong to the SR proteins, RNA-binding motif (RBM) proteins, et al. An article with research on the mechanism of hnRNPs is the eighth cited reference (**Table 4**) and also ranks fifth in the strength of the reference burst (**Figure 6C**).

In addition to a carboxy-terminal arginine/serine-rich domain that contributes to protein-protein interactions, SR proteins have one or two copies of an RNA recognition motif domain at the amino terminus that offers RNA-binding specificity(111,112). The majority of SR proteins function as splicing activators, helping the spliceosome recognize exons and enable exon inclusion by binding to pre-mRNA at exonic splicing enhancers. SR proteins frequently face competition from splicing repressors such as hnRNPs. By binding to exonic or intronic splicing silencers, hnRNPs obstruct spliceosome elements’ access and suppress splice site choice. RNA-binding domains and somewhat unstructured domains, which are likely involved in protein-protein interactions, are both present in hnRNPs in a comparable manner. Exon skipping is prevented by SR proteins’ concentration-dependent inhibition of hnRNPs activity(110,113,114)

In conclusion, many RNA-binding proteins, such as SR proteins and hnRNPs, bind splicing enhancers and silencers(115). Besides, some members of RBM proteins also play important physiological roles as splicing factors. Similar to our study, among them, there are more reports about RBM4(116), RBM5(117), and RBM10(118). Similar to SR protein and hnRNPs protein, RBMs can also regulate the occurrence of splicing events alone or cooperate with other splicing factors to regulate the splicing process(119).

### 4.3 Strength and Limitations

Overall, as far as we know, this article may be the first to apply bibliometric methods to comprehensively examine papers associated with AS of cancer research published in the last ten years. The bibliometric method offers a fresh and unbiased perspective on the changing research hotspots and fashion in contrast to conventional reviews(21). Simultaneously, we conducted an investigation using various bibliometric tools, which could produce more prosperous outcomes across numerous dimensions(21,120). This study will educate the public about the significance of AS of cancer, offer scholars a complete image of AS of cancer study, and additionally provide a detailed and impartial direction for the field’s upcoming growth. This study unavoidably has certain shortcomings. First, we only obtained the WoSCC database’s English-language articles, leaving out non-English or non-WoSCC items. However, WoSCC’s English articles are the most often utilized data source in bibliometrics, thus to some extent they may represent the majority of the field(21,121). Secondly, Some studies report that bibliometric methods are inevitably biased because they are based on natural language processing(17,20). Our findings, however, are comparable to recent conventional reviews while offering more comprehensive and unbiased data(4,11,80).

## 5 CONCLUSION

In conclusion, research on AS of cancer has steadily advanced step by step with active cooperation over the past ten years, with the possibility that the United States will continue to hold the lead in this field. Scorilas Andreas and Wang Eric T. were the authors with the most publications and co-citations in AS of the cancer field, respectively. Currently, AS of cancer research is mainly centred on hallmarks of AS in cancer, AS signatures and therapeutic targets, and further mechanisms. Among them, splicing factors which regulate EMT or other hallmarks, further aberrant AS signatures and therapeutic targets in cancer research, and the mechanism of AS may become popular and fruitful directions. These could offer direction and fresh perspective for future studies in the AS of cancer.

## ACKNOWLEDGEMENTS

Not applicable.

## DATA AVAILABILITY STATEMENT

The original contributions presented in the study are included in the article/Supplementary Material. Further inquiries can be directed to the corresponding authors.

## AUTHOR CONTRIBUTIONS

WLW, PYN, TB, BY and LZS designed this study. TB, BY, BDJ and GY collected the data and performed the analysis. ZX, ZSW and ZYH provided support in the data curation, validation and visualization. TB, BY, and BDJ wrote the original draft. WLW and LZS reviewed and revised the manuscript. All authors reviewed the manuscript.

